# Type 2 Deiodinase in Cancer-Associated Fibroblasts is required to sustain growth of poorly and undifferentiated thyroid cancer

**DOI:** 10.1101/2025.07.10.664131

**Authors:** Maria Angela De Stefano, Cristina Luongo, Tommaso Porcelli, Costantina Cervone, Stefano Spiezia, Claudia Misso, Vincenza Cerbone, Anna Maria Carillo, Giancarlo Troncone, Martin Schlumberger, Domenico Salvatore

## Abstract

Poorly differentiated (PDTC) and anaplastic thyroid carcinomas (ATC) are characterized by rapid progression and poor patient survival. While the tumor microenvironment (TME) - particularly cancer-associated fibroblasts (CAFs)- plays a crucial role in supporting tumor growth, its metabolic contribution remains poorly understood. Here, we identify a critical role for type 2 deiodinase (D2), the enzyme that activates the thyroid hormone (TH) thyroxine (T4) into the biologically active triiodothyronine (T3), in sustaining a pro-tumorigenic TME in PDTC and ATC. We show that D2 is expressed not only in thyroid epithelial cancer cells, but at even higher levels in CAFs, especially inflammatory CAFs (iCAFs). In *in vivo* mouse models, pharmacological inhibition of D2 reduces tumor growth and alters composition of CAFs. In 3D co-culture spheroids, D2 activity proves essential for supporting tumor cell proliferation by establishing a paracrine loop between stromal and epithelial cancer cells that amplifies local TH signaling. Notably, human PDTC organoids expressing D2 respond to modulation of TH levels, confirming the functional relevance of this metabolic axis in human tumors. In conclusion, these findings identify D2 as a key mediator of stromal-epithelial crosstalk in PDTC and ATC, and highlight local TH metabolism as a potential therapeutic target in these lethal cancers.

## INTRODUCTION

While most thyroid cancers are well-differentiated and characterized by a low proliferation rate —such as papillary thyroid carcinoma (PTC) and follicular thyroid carcinoma (FTC)— these tumors can progress to more aggressive forms, including poorly differentiated thyroid carcinoma (PDTC) and anaplastic thyroid carcinoma (ATC), which exhibit a rapid cell proliferation, local invasion, and early metastatic dissemination (1, 2)

Beyond intrinsic properties of the epithelial thyroid cancer cells, the tumor microenvironment (TME) has emerged as a key driver in PDTC/ATC progression (3–5). Among its stromal components, cancer-associated fibroblasts (CAFs) actively support tumor growth, immune evasion, angiogenesis, and extracellular matrix (ECM) remodeling. Importantly, CAFs are a heterogeneous population: myofibroblastic CAFs (myCAFs) reside in close proximity to tumor cells, contribute to ECM remodeling, and exhibit high contractility; inflammatory CAFs (iCAFs) are more dispersed and secrete pro-inflammatory cytokines like IL-6, IL-1β, CXCL1/2, and LIF; antigen-presenting CAFs can modulate immune responses by interacting with CD4+ T cells through MHC class II presentation (6, 7). In ATC, CAFs are particularly abundant and predominantly exhibit the highly activated phenotype of iCAF, (8), though their molecular interactions with epithelial tumor cells remain largely unexplored.

An emerging and underappreciated aspect of the TME involves regulation of local, intra-tumoral thyroid hormone (TH) metabolism and signaling. Thyroid hormones—thyroxine (T4) and its biologically active form triiodothyronine (T3)—are critical regulators of metabolism, proliferation, and differentiation (9). The local conversion of T4 to T3 is catalyzed by type 2 deiodinase (D2) enzyme, which is expressed in specific cells – including fibroadipogenic progenitors –in selected tissues and several cancers (10, 11). In contrast, type 3 deiodinase (D3) inactivates T4 and T3, reducing local TH availability (12, 13). The balance between D2 and D3 expression governs the intracellular T3 concentration, and this is their only known function (14).

We and others have shown that early-stage epithelial tumors, such as basal cell carcinoma (15) and colon cancer (16) display high D3 and low D2 expression leading to localized intracellular hypothyroidism. As these tumors progress, D3 level typically declines and D2 expression increases, suggesting a dynamic metabolic reprogramming that favors T3-driven tumor growth at advanced stages (16).

Similarly, in differentiated thyroid cancers, such as PTC, D3 is frequently upregulated - especially in BRAF-mutated tumors (17), while D2 is suppressed versus healthy thyroid tissue, likely contributing to an intracellular hypothyroid microenvironment. In contrast, immortalized human ATC cells exhibit high D2 expression and we have previously shown that D2 promotes their survival (18). Noticeably, we have also demonstrated that TP53 inactivation, a hallmark of many PDTC and ATCs but absent in differentiated thyroid cancers, enhances D2 expression in advanced thyroid and skin cancers (18, 19, 20).

In this study, we investigate the role of D2 in PDTC/ATC tumorigenesis, with a specific focus on its expression and function in both epithelial tumor cells and CAFs. By using mouse models and human patients derived-organoids, we show that D2 is expressed not only in the epithelial tumor cells but also - and even at higher levels - in the mesenchymal compartment. We propose that D2-dependent activation of local thyroid hormone signaling sustains the aggressive phenotype of a subset of PDTC/ATCs, representing a novel metabolic axis within the TME that could be therapeutically exploited.

## RESULTS

### D2 expression dynamics and cellular distribution during thyroid cancer progression

To get insight on the expression of deiodinases among distinct cell populations in undifferentiated tumors, we analyzed their expression profile during tumor progression using a well-characterized mouse model of ATC (TPO-Cre/LSL-BrafV600E/p53^lox-lox^/eYFP (21, 22) (Figure 1A). At early stages of tumorigenesis (6–8 weeks), when thyroid glands reach approximately 50 mg (about 10 times their normal weight), histology resembles papillary thyroid carcinoma (PTC) (Figure 1B-D) and D3 is the predominantly expressed deiodinase (Figure 1E). By 18–20 weeks, when thyroid weight exceeds 100 mg and histology is consistent with PDTC/ATC, D2 expression increases markedly, while D3 level declines (Figure 1B–E). This reciprocal expression pattern indicates that D3 and D2 are dynamically and inversely regulated during thyroid tumor dedifferentiation and progression.

**Figure 1.**
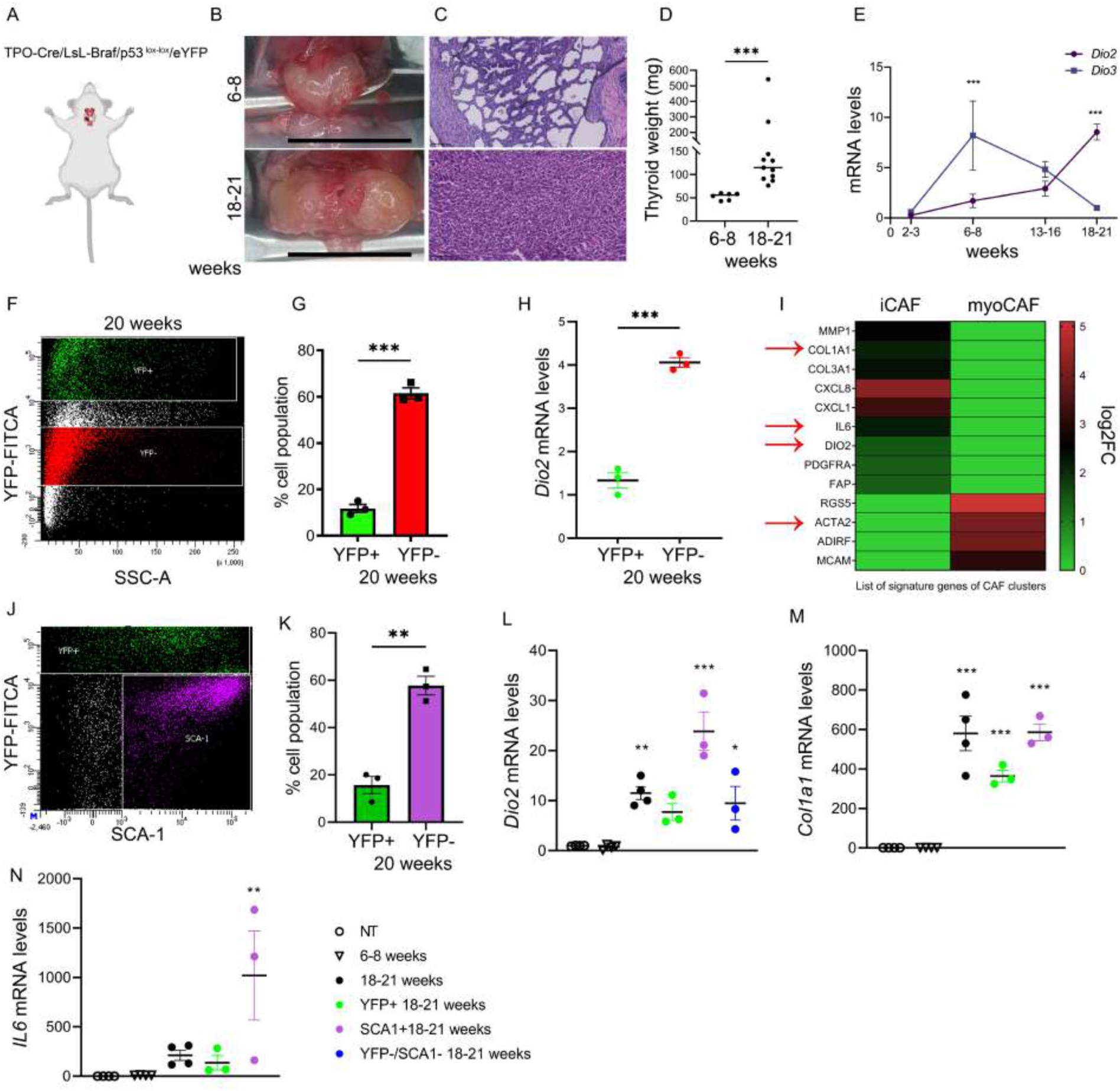
D2 is expressed in distinct cell population of thyroid tumors. (A) Thyroid cancer mouse model: TPO-Cre/LSL-BrafV600E/p53^lox-lox^/eYFP. (B) Representative images of thyroid tumors in the ATC mouse model at 6-8 and 18-21 weeks of age (Scale bar, 1cm). (C) Images of H&E staining of thyroid tumor section from ATC mouse model at 6-8 and 18-21 weeks (Scale bar, 100 *μ*m). (D) Tumor thyroid weights from ATC mice at 6-8 and 18-21 weeks. Each dot represents an individual animal (6-8 weeks, n=6; 18-21 weeks, n=11). (E) D2 and D3 mRNA expression levels in thyroids from ATC mice at different ages compared to normal thyroid tissue. Each point represents the mean of pooled samples from several animals (2-3 weeks, n= 3; 6-8 weeks, n=6; 13-15 weeks, n=7; and 18-21 weeks, n= 11). (F) Representative flow cytometry dot plot showing YFP^+^ (green) and YFP^−^ (red) cells. (G) Percentage of epithelial (YFP^+^) and mesenchymal (YFP^−^) cells from pooled thyroid tumors of ATC mice at 20 weeks, analyzed by FACS. Each point represents an individual experiment (pooled tumors from n=3 mice). (H) D2 mRNA expression in YFP^+^ and YFP^−^ cells sorted from ATC thyroid tumors at 20 weeks. Each point represents a technical replicate from pooled tumors (n=3 mice). (I) In silico analysis of scRNA-seq data extracted from Lu et al. (23). (J) Representative flow cytometry dot plot showing YFP^+^ (green) and SCA1^+^ (violet) cells. (K) Percentage of epithelial (YFP^+^) and CAF (SCA^+^) cells in pooled thyroid tumors from 20-week-old ATC mice, analyzed by FACS. Each point represents an individual experiment (pooled tumors from n=3 mice). (L) D2 mRNA expression in thyroid tumors at 8 and 20 weeks and in YFP^+^ and SCA^+^ and YFP^−^/SCA1^−^ cells sorted at 20 weeks. (M-N) Col1a1 (M) and IL6 (N) mRNA expression in thyroid tumors at 8 and 20 weeks and in YFP^+^ and SCA^+^ cells sorted from tumors at 20 weeks. Each point represents an individual experiment. Data are expressed as mean ± S.E.M. * P < 0.05, ** P < 0.01, ***P < 0.0001 using t tests in D, G, H and K or ANOVA in E, L, M and N.

Given the marked expression of D2 in advanced-stage thyroid tumors, we analyzed its cellular source by separating epithelial (YFP⁺) and non-epithelial (YFP⁻) cells from 20-week-old thyroid tumors using fluorescence-activated cell sorting (FACS) (Figure 1F). YFP⁺ cells we selected accounted for ∼15%, while YFP⁻ for ∼60% of the total viable cell population (Figure 1G). *Dio*2 mRNA was detected in both cell compartments, but was about threefold higher in the YFP⁻ fraction (Figure 1H), indicating that D2 is predominantly expressed in non-epithelial cells within undifferentiated tumors.

This observation aligns with analyses of single-cell RNA-sequencing datasets from human ATC (publicly available), which show high D2 expression in cancer-associated fibroblasts (CAFs)—particularly in the inflammatory CAFs (iCAFs) subtype, but minimal expression in myofibroblastic CAFs (myCAFs) (Figure 1I) (23). The *Dio*2 expressed in mesenchymal cells is also consistent with our prior identification of *Dio*2 expression in fibroadipogenic progenitors isolated from mouse healthy limb muscles (FAPs) (24), a stromal cell population which is a source of fibroblasts and of CAFs, hereafter referred to as preCAFs cells (25, 7).

To further identify the specific non-epithelial (YFP⁻) cell population that contributes to D2 expression, we conducted additional FACS analyses to isolate Sca1⁺ cells (a general marker of CAFs) alongside YFP⁺ epithelial cells from 20 weeks-old tumors (Figure 1J). At this stage, Sca1⁺ CAF population accounted for ∼60% of viable cells (Figure 1K), a significant increase from the ∼30% of SCA1^+^ CAF cells identified at 6-8 weeks (PTC-like phase, Supplemental Fig. 1A, B) indicating substantial remodeling of the tumor microenvironment. *Dio*2 expression was significantly enriched in the Sca1⁺ CAF population at 20 weeks, with levels approximately three times higher than in epithelial cells (Figure 1L). These CAFs were likely iCAF, as indicated by high expression of iCAF specific markers (Col1a1 and IL6, see Figure 1I, M, N) and the near absence of ACTA2 expression, a marker of myCAF, which encodes for the αSMA protein (Figure 1I and Supplemental Fig. 1C). Notably, both iCAF cell markers were virtually undetectable at the earlier (6-8 weeks) stage as well as in the healthy thyroid gland, consistent with the tumor stroma remodeling and the shift in deiodinases expression observed during tumor progression (Figure 1L-N).

Taken together, these data show that D2 is highly expressed in CAFs and in particular in a specific CAF subpopulation, inflammatory CAFs (iCAFs). To explore whether other non-epithelial/non-CAF cells (including TAMs and CD45⁺ leukocytes) also express D2, we analyzed the YFP⁻/ Sca1⁻ residual population at 20 weeks. *Dio*2 was also detected at levels roughly 2.5 times higher than in epithelial tumor cells (Figure 1L), suggesting additional contributors to intra-tumoral D2 expression.

Overall, these results indicate that D2 is expressed in ATC in multiple cell compartments, and at the highest levels in iCAFs. The expansion of CAFs—especially iCAFs—correlates with a specifically tuned thyroid hormone metabolism, wherein D2 is the dominant expressed deiodinase.

### D2 inhibition alters the composition of CAFs in ATC xenograft tumors

To investigate the role of D2 in both epithelial and mesenchymal components of anaplastic thyroid cancer (ATC) *in vivo*, we performed xenograft experiments using human ATC 8505 cells, which express D2 (18). Cells were subcutaneously injected into nude mice, and tumors were monitored and harvested 8 weeks post-inoculation (Figure 2A). To distinguish human *DIO*2 expression (originating from the human 8505 cells) from potential host-derived murine *Dio*2 expression, we designed species-specific oligonucleotides (Supplemental Fig. 2A, B).

**Figure 2.**
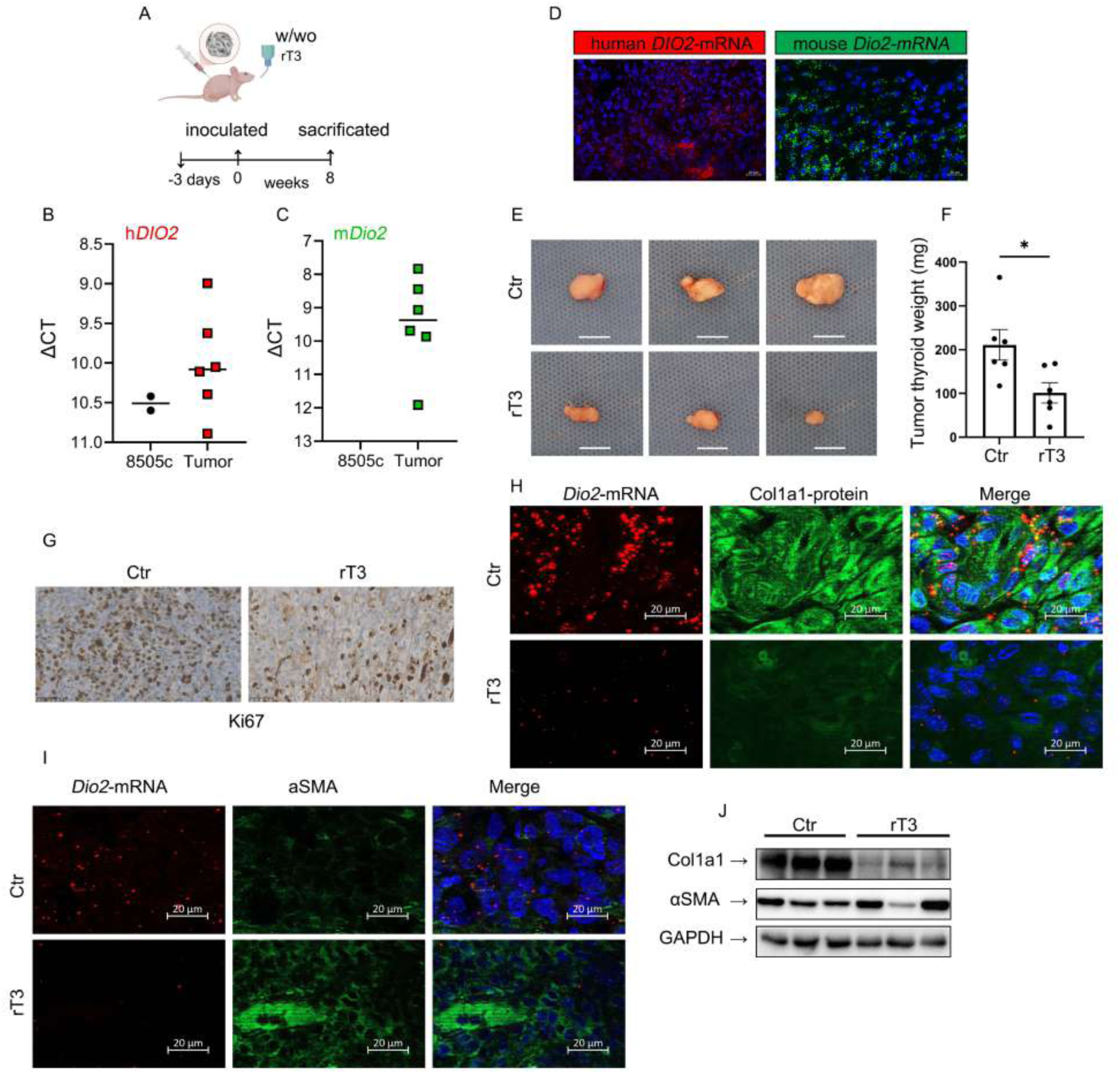
Effects of D2 inhibition on xenograft thyroid tumors. (A) Experimental design: 8505 cells were injected into nude mice treated or not with rT3 for three days prior to inoculation and for an additional eight weeks. (B-C) Relative expression (ΔCT values) of human (B) and mouse (C) D2 in formed tumors, using species-specific oligonucleotides. Non-injected 8505 cells were used as positive control for human and negative control for mouse probes. Each point represents an individual animal. (D) Representative RNAscope fluorescent images showing single mRNA molecules of human *DIO*2 (red dots) and mouse *Dio*2 (green dots). DAPI (blue) marks nuclei (Scale bar, 20 *μ*m). (E) Representative tumor images excised from rT3-treated and control nude mice (Scale bar, 1cm). (F) Tumor weights from rT3-treated and control groups. Each dot represents an individual animal (control n=6; rT3-treated n=5). (G) Ki67immunohistochemistry of tumors from rT3-treated and control groups (Scale bar, 50 *μ*m). (H-I) Representative dual RNAscope for Dio2 (red) and protein staining for Col1a1 (H) or αSMA (I) in tumors from the rT3-treated and control groups (Scale bar, 20 *μ*m). (J) Western blot analysis of Col1a1 and αSMA protein levels in tumors from rT3-treated and control mice (n = 3 per group); Each lane represents an individual animal. Data are expressed as mean ± S.E.M. * P < 0.05, ** P < 0.01, ***P < 0.0001 using t tests.

Firstly, we observed that human *DIO*2 expression is maintained in ATC tumors *in vivo* compared to the *in vitro* levels in 8505 cells, but with a much higher variability *in vivo* (Figure 2B). Notably, mouse *Dio*2 mRNA was also abundantly expressed in the xenograft tumors (Figure 2C), suggesting recruitment of host-derived stromal cells expressing *Dio*2. By RNAscope analysis, we found that human and murine D2 mRNA showed different localization patterns within the tumors. Human D2 mRNA was enriched in the central tumor region, whereas mouse D2 mRNA was prominent at the tumor periphery (Figure 2D and Supplemental Fig. 2C). These results confirmed that both tumor-derived epithelial cells and host-derived stromal cells contribute to intra-tumoral D2 expression.

To explore the functional role of D2 *in vivo*, we treated mice with orally administered reverse-T3 (rT3), a known enzymatic inhibitor of D2, with no activity at the thyroid hormone receptor level (Figure 2A). After 8 weeks, mice were sacrificed and tumors collected. Tumors from rT3-treated mice were significantly smaller than those from vehicle-treated controls (Figure 2E, F), indicating that D2 promotes ATC growth. Untreated tumors exhibited classic high-grade PDTC/ATC features, including nuclear pleomorphism, high mitotic rate, necrotic areas and disorganized architecture (Figure 2G and Supplemental Fig. 2D-H). Tumors from D2-inhibited rT3 treated mice appeared more organized, with reduced nuclear pleomorphism, mitotic rates (Supplemental Fig. 2D, E) and reduced Ki-67 staining (Figure 2G and Supplemental Fig. 2F), indicating a decreased proliferation rate.

Next, we analyzed the role of D2 in determining the composition of CAFs in ATC xenograft tumors. By using a dual RNAscope/immunofluorescence (IF) staining, we found that mouse *Dio*2 mRNA signal was mostly localized with Col1a1 protein staining (a marker of iCAF) (Figure 2H). Interestingly, D2-inhibition with rT3 resulted in decreased Col1a1 mRNA and protein expressions, while aSMA (which marks the myCAF) was largely unaffected (Figure 2H-J).

Collectively, these findings indicate that D2 expression marks the iCAF, but not myCAFs, population within the ATC tumor. Blocking D2 impairs tumor growth and selectively alters CAF composition by reducing the iCAFs.

### Role of D2 in CAFs in heterotypic spheroids

To get insight in the functional role of D2 in CAFs, we established heterotypic 3D spheroid cultures composed of human ATC 8505 cells and mouse primary fibroadipogenic progenitors freshly isolated from healthy limb muscle (preCAFs, i.e. CAF precursors (26) at a 1:3 ratio (see Material & Methods and Supplemental Fig. 3A, B). This 3D cell system was mandatory as D2 expression was completely lost after 7 days in preCAFs kept in 2D monolayer culture (Supplemental Fig. 3C and (27)).

When heterotypic spheroids (8505 cells/preCAFs) were kept in 3D culture, they exhibited a significantly enhanced linear growth compared to mono-type spheroids consisting of ATC 8505 cells (Supplemental Fig 3D) indicating growth cooperative interactions in the spheroids between epithelial and mesenchymal cells. Moreover, treatment with rT3 significantly reduced the growth of heterotypic 8505/preCAFs spheroids compared to the mono-type 8505 spheroids (Figure 3A-C). Notably, mono-type preCAF spheroids showed a non-significant proliferation, which was not affected by rT3 (Figure 3A).

**Figure 3.**
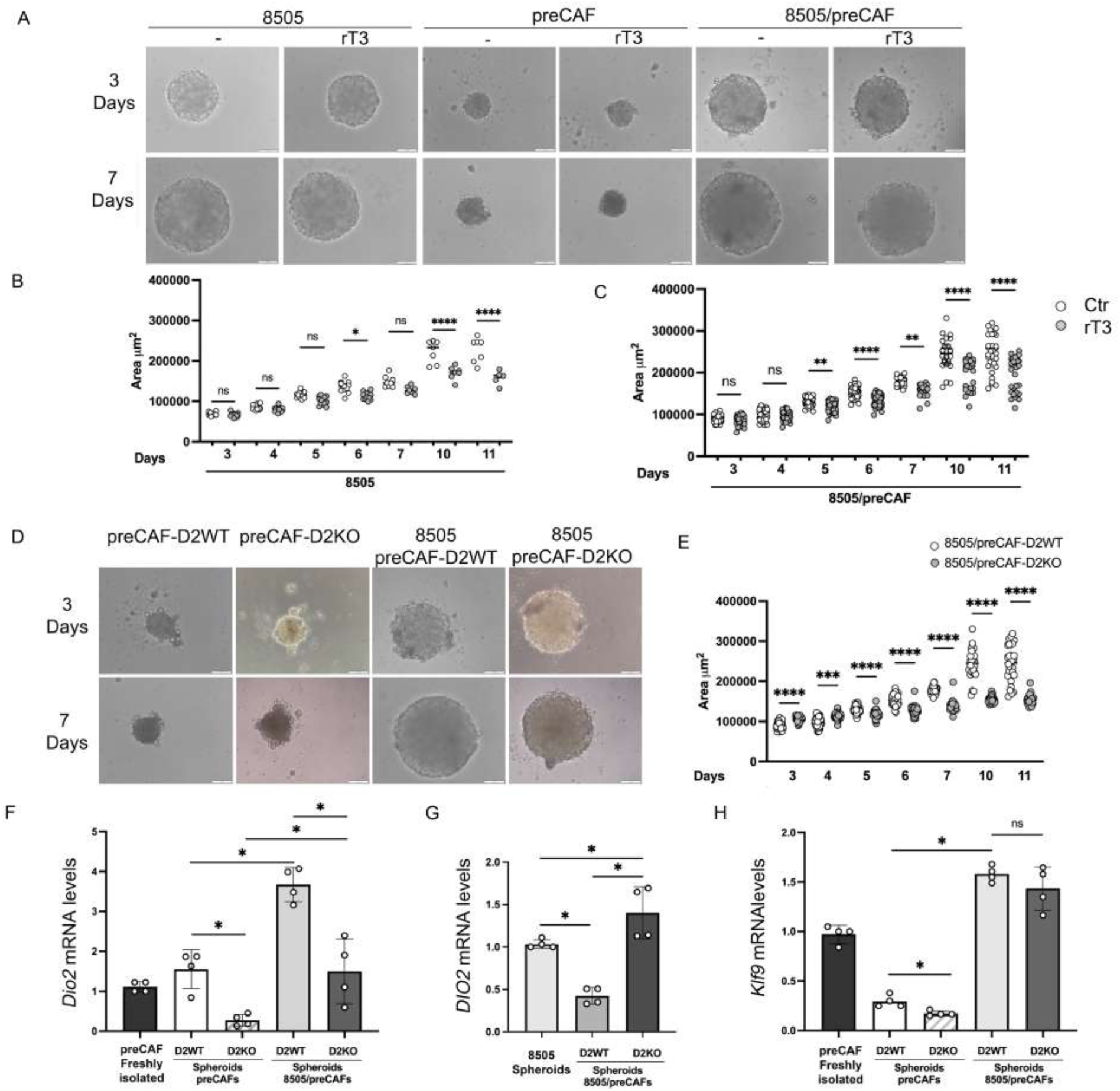
Role of CAF-D2 in heterotypic spheroids. (A) Representative brightfield (BF) images of mono-type (8505 or preCAF) and hetero-type (8505/preCAF) spheroids cultured for 3 and 7 days with or without 30 nM rT3 (Scale bar, 100 *μ*m). (B-C) Time-course analysis of spheroid area in mono-type 8505 (B) and hetero-type 8505/preCAF (C) spheroids cultured for 3 and 7 days in the presence or absence of rT3. D) Representative BF images of mono-type preCAF-D2WT and preCAF-D2KO spheroids, and hetero-type 8505/preCAF-D2WT and 8505/preCAF-D2KO spheroids (Scale bar, 100 *μ*m). (E) Time-course analysis of spheroid area for hetero-type 8505/preCAF-D2WT and 8505/preCAF-D2KO spheroids. (F) Mouse *Dio*2 mRNA levels in freshly isolated preCAFs, mono-type preCAF-D2WT and preCAF-D2KO spheroids, and hetero-type 8505/preCAF-D2WT and 8505/preCAF-D2KO spheroids at 7 days post-seeding. (G) Human *DIO2* mRNA levels in mono-type 8505 spheroids and hetero-type 8505/preCAF-D2WT and 8505/preCAF-D2KO spheroids at day 7 post-seeding H) Mouse *Klf9* mRNA levels of mouse in freshly isolated preCAFs, mono-type preCAF-D2WT and preCAF-D2KO spheroids, and hetero-type 8505/preCAF-D2WT and 8505/preCAF-D2KO spheroids at 7 days post-seeding. Each dot represents an individual spheroid. Data are expressed as mean ± S.E.M. * P < 0.05, ** P < 0.01, ***P < 0.001, ****P < 0.0001 using ANOVA test.

To specifically assess the role of D2 expressed in the preCAFs population in the spheroid’s growth, we generated heterotypic spheroids using ATC 8505 cells and either D2 wild-type (preCAF-D2WT, used in the previous experiments) or D2 knock-out preCAFs (preCAF-D2KO), in which *Dio*2 was genetically deleted *ex-vivo* in freshly isolated preCAFs (see Materials & Methods). Consistent with the pharmacological D2 inhibition with rT3 (Figure 3A), *Dio*2 genetic deletion in preCAFs did not affect their own growth in mono-type spheroids (Figure 3D). However, heterotypic spheroids containing ATC 8505/preCAF-D2KO cells grew significantly less than those with preCAF-D2WT, demonstrating that the cell-specific deletion of D2 in CAFs (with normal D2 in ATC 8505 cells) reduces tumor cell proliferation and overall spheroid expansion (Figure 3D, E).

We next examined the regulation of human and mouse D2 expression in co-cultured spheroids (Figure 3F, G). In ATC 8505/preCAF-D2WT spheroids, mouse *Dio*2 expression was significantly upregulated in preCAFs upon co-culture with ATC 8505 cells, compared to preCAF-D2WT cultured alone (Figure 3F). This induction was functionally relevant, as shown by the increased expression of mouse *Klf9*, a well-known thyroid hormone (TH)-responsive gene (Figure 3H), suggesting that preCAFs acquired enhanced TH signaling upon interaction with tumor cells.

Interestingly, human *DIO*2 expression in ATC 8505 cells was reduced in the presence of D2-expressing preCAFs (Figure 3G). This down-regulation was not observed in spheroids with preCAF-D2KO cells, supporting the hypothesis of a reciprocal D2 balance in the tumor microenvironment (Figure 3F–H).

Overall, these data suggest that a finely tuned D2 expression between ATC 8505 and preCAF is part of the cross-talk between the epithelial and stromal components, which is necessary for tumor growth.

### Effects of D2 inhibition on human PDTC/ATC-derived organoids

Having established a role for D2 in promoting ATC growth through epithelial–mesenchymal interactions in experimental models, we sought to determine whether similar mechanisms operate in human tumors. To this end, we analyzed *DIO*2 mRNA expression in three consecutive advanced thyroid cancers, including two poorly differentiated and one papillary, surgically resected at our institution (Figure 4A, Table S1 and Supplemental Fig. 4). Among these tumors, *DIO*2 was detected in the 2 PDTCs (#1 and #2), with an expression level exceeding that of healthy thyroid tissue used as a reference (Supplemental Figure 4A), while absent in the PTC. The expression of the other deiodinases (*DIO*1 and *DIO*3) was extremely variable among the tumor samples (Supplemental Fig. 4 B, C), while thyroid-differentiation markers TG, TPO, PAX8, NIS and TTF1 (also known as NKX2.1) were reduced in all tumors, compared to healthy thyroid tissue (Supplemental Fig. 4 D, E). The presence of *DIO*2 mRNA was further validated by RNAscope, which showed specific *DIO*2 mRNA signals in *DIO*2-positive tumors (Supplemental Fig. 4 I). The expression of D2 in PDTC samples is in agreement with *in silico* data showing the highest expression of *DIO*2 mRNA in PDTCs while absent in PTCs (Supplemental Fig. 4J).

**Figure 4.**
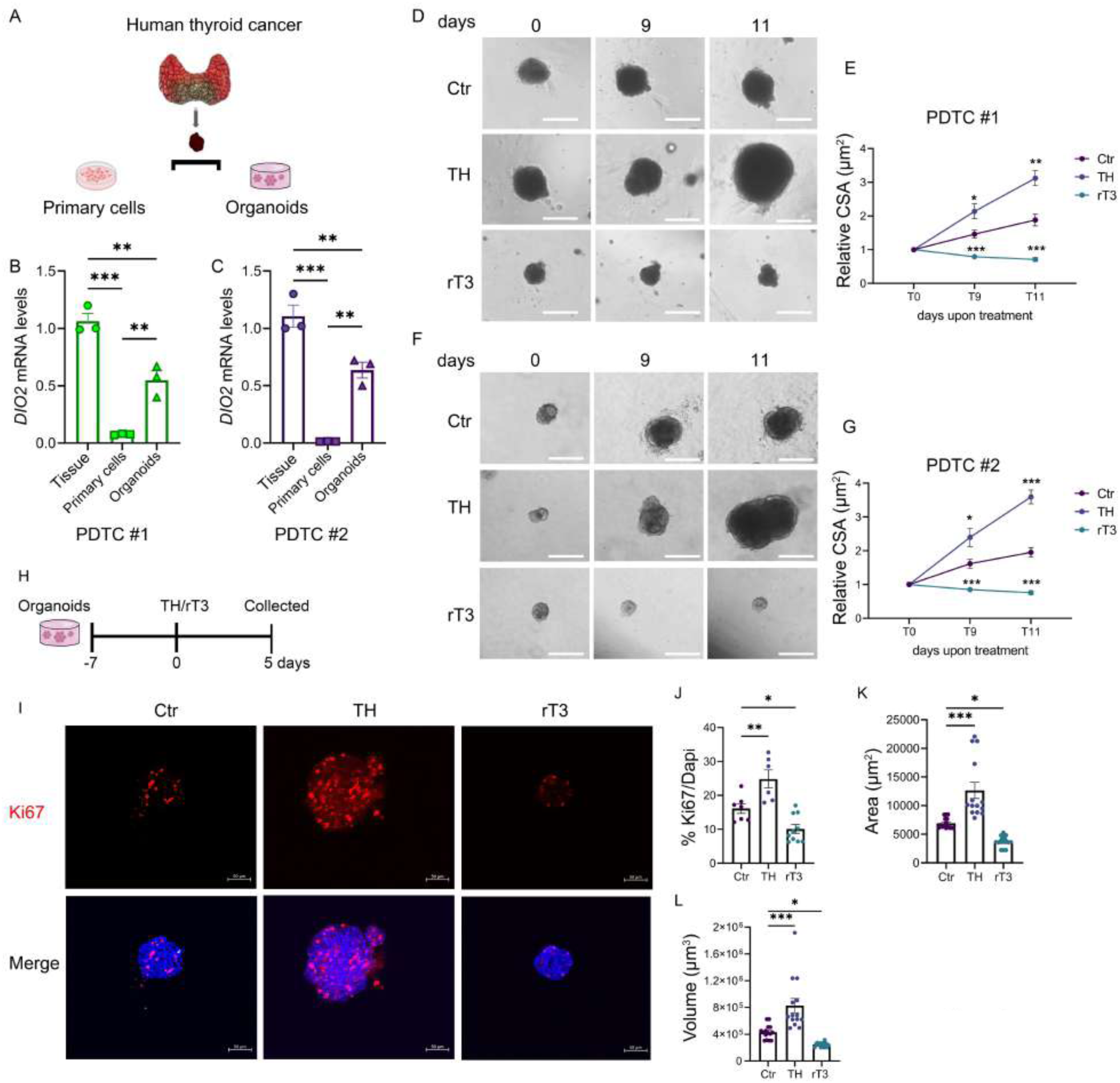
Effects of D2 inhibition on human PDTC-derived organoids. (A) Schematic of derivation of primary cells and organoids from patient tumor tissues. (B-C) DIO2 mRNA expression in tumor tissues, derived primary cells, and derived organoids from patients PDTC #1 (B) and PDTC #2 (C). Each point represents a technical replicate. (D-E) Representative BF images of patient-derived organoids (patients PDTC #1 (D) and relative cross-sectional area (E), upon treatment with TH and rT3 at different time points (Scale bar, 200 µm), areas were normalized to their respective day 0 area of treatment. (F-G) Representative BF images of patient-derived organoids (patients PDTC #2 (D) and relative cross-sectional area (E), upon treatment with TH and rT3 at different time points (Scale bar, 200 µm), areas were normalized to their respective day 0 area of treatment. (H) Schematic of organoid experiment (patient PDTC #1) treated with TH and rT3 for 5 days. (I) Representative Ki67 immunofluorescence of organoids. (J) Quantification of Ki67^+^ cells normalized to DAPI. (K-L) Quantification of organoid area in *μ*m^2^ (K) and volume in *μ*m^3^ (L) of experiment in H. Each point represents an individual organoid. Data are expressed as mean ± S.E.M. * P < 0.05, ** P < 0.01, ***P < 0.0001 using ANOVA test.

To assess D2 levels in organoids and cell cultures compared to primary tumor, each tumor sample was split into two portions: one for conventional two-dimensional (2D) primary culture and the other for three-dimensional (3D) organoid culture (Figure 4A). Organoids formed after ∼1 week and were maintained for 2–3 months. In *DIO*2-positive tumors (PDTC # 1 and PDTC #2), we measured *DIO*2 mRNA in both 2D and 3D cultures and compared it to the original bulk tissue. After 10 days, *DIO*2 expression was almost completely lost in 2D cultures, but partially retained in 3D organoids (Figure 4B, C) and remained stable thereafter (data not shown).

To test the impact of thyroid hormone level alterations, organoids from the 2 PDTCs were treated with either exogenous TH (mimicking a hyperthyroid state) or rT3 (lowering intracellular T3 levels by D2 inhibition) and compared to untreated controls. Organoids growth increased in response to TH, while rT3 treatment significantly reduced their expansion (Figure 4D-G). Ki67 staining revealed a corresponding increase in proliferative cells in T3-treated organoids, and a decrease following rT3 exposure (Figure 4H, I). Consistent with these findings, organoids area and volume were significantly increased with TH treatment and reduced or unchanged following rT3 treatment (Figure 4J, K). In contrast, treatment with rT3 had no significant effect on the growth of the organoids derived from the PTC that did not express D2 (Supplemental Fig 4K, L).

Altogether, these data indicate that *DIO*2 that is expressed in human poorly differentiated thyroid cancers plays a functional role in promoting their growth through local T3 production. The observed responsiveness of these human cancer-derived organoids to both exogenous TH and D2 inhibition further supports the relevance of thyroid hormone signaling in thyroid cancer progression.

## DISCUSSION

In this study, we demonstrate that D2 is expressed in both mouse and human PDTC and ATC, with localization in epithelial as well as stromal cells, particularly within the CAF population. In CAFs, D2 is a critical metabolic player that controls tumor growth and CAF’s identity by sustaining local TH activation. PDTC/ATCs no longer express D3, which marks thyroid tumor cells at early (PTC-like) stage of tumor formation (here and (28).

It is important to note that the only known biological function for D2 enzyme is to convert T4 into T3, the most active thyroid hormone. Thus, D2 expression typically indicates increased intracellular T3 concentration. This is consistent with the observed upregulation of *Klf9* - a well-characterized thyroid hormone responsive gene (29)-in CAFs, upon increased D2 expression (see Figure 3). Notably, D2 is a highly efficient yet unstable enzyme, with a very short half-life of less than 30 minutes. This likely explains the failure so far to develop effective anti-D2 antibodies needed to monitor D2 protein, despite the substantial efforts of several research groups, including ours. However, D2 mRNA is readily measurable and accurately reflects enzyme activity (30); therefore, we used D2 mRNA levels as a proxy for D2 expression and enzymatic activity. The reciprocal expression profile of D3 and D2 during tumor progression suggests that the D3/D2 ratio might serve as a future biomarker for thyroid cancer grading. Altered expression or activity of D2 has been implicated in various cancer types, with complex and sometimes contradictory roles depending on the tumor type and microenvironment (31). One of the best-characterized tissues concerning the expression of deiodinases is the skin, particularly regarding the difference from the non-aggressive basal cell carcinoma (BCC) and the more invasive squamous cell carcinoma (SCC). In BCC, D2 is not present, while D3 is upregulated, suggesting a suppression of T3 signaling. In contrast, SCC exhibited strong D2 expression, induced by mutant p53. In this highly aggressive setting, genetic deletion of *Dio*2 significantly impairs tumor progression and reduces epithelial– mesenchymal transition (EMT) (20). We have previously shown that D2 is downregulated in papillary thyroid carcinoma (PTC) but upregulated in the more aggressive PDTC/ATC (11). This expression pattern closely parallels what was observed in colorectal cancer, where D2 is suppressed in early stages but becomes upregulated in advanced, aggressive tumors, enhancing intracellular T3 availability (16).

Cancer-associated fibroblasts (CAFs) are pivotal components of the tumor microenvironment (TME). They exert a profound influence on cancer cell behavior and actively contribute to tumor progression through complex interactions with cancer cells and other stromal constituents. Due to their inherent plasticity, CAFs can be “educated” by cancer cells, leading to dynamic, context-dependent changes in their phenotype and function. The pre-CAF population we utilized in the present study constitutes a major source of this mesenchymal component. These cells, being progenitor-like, exhibit a heightened capacity to receive and integrate multiple signaling cues from the tumor microenvironment.

In ATC, the TME is notably enriched with CAFs, particularly the inflammatory subtype (iCAFs)—the most “activated” CAF population—which replaces the myofibroblastic CAFs (myCAFs) commonly found in less aggressive forms of thyroid cancer (23). iCAFs are characterized by the expression of numerous inflammation-related genes, including CXCL1, CXCL6, and CXCL8 as well as COL1A1, COL1A2, and COL3A1, which encode essential structural components of the extracellular matrix (ECM). Importantly, CAFs also express thyroid hormone transporters such as MCT8 and MCT10, as well as deiodinases, particularly D2 and D3 (24), allowing them to finely regulate local TH availability within the TME. These features align with the broader actions of TH signaling, which modulates genes involved in ECM remodeling, growth factor signaling, inflammation, and immune evasion (9).

Our findings demonstrate that D2 is highly expressed in iCAFs in PDTC/ATC, underscoring the potential requirement for elevated local T3 levels in shaping the thyroid TME. Notably, behind CAF, we also detected D2 expression in the excluded fractions from the FACS sorting (i.e. the cells negative for YFP and SCA1) (Figure 1). Stromal cells are rare in well differentiated PTC and FTC, but are present with an increasing number and related to worst prognosis in PDTC and in ATC (32). This cell fraction includes blood-derived cells, such as macrophages, which potentially represent up to 50% of total cell population in human ATCs (33). In non-cancerous contexts, D2 is known to be highly expressed in macrophages, particularly during their M2 polarization (34, 35), a phenotype that predominates in ATCs (36). The role of D2 in cancer macrophages was not addressed in this study and indeed warrants further investigations.

Using 3D spheroid cultures and *in vivo* models, we showed that D2 activity in CAFs is essential for sustaining tumor spheroids growth (Figure 3). Interestingly, D2-deficient CAFs failed to support spheroids growth as effectively as the wild type cells (Figure 3), indicating that D2 contributes to a paracrine-signaling environment that favors tumor growth. This is further highlighted by the observed balance in D2 expression between epithelial and mesenchymal compartments, measured in normal conditions and upon D2 genetic deletion (Figure 3E-G).

Beyond supporting epithelial proliferation, D2 also drives the phenotypic programming of the TME, controlling CAF fate. In xenograft models, T3 production via D2 was identified as a critical determinant of iCAF identity and maturation, with functional consequences for tumor progression. Inhibition of D2 impairs the expression of iCAF markers (e.g., Col1a1, IL6), while the expression of myCAF markers (e.g., αSMA) remained unaffected (Figure 2). These findings suggest the existence of a paracrine-signaling loop within the TME wherein thyroid tumor cells – stimulated by potent oncogenic drivers such as mutated p53 gene, while repressing D3, increase D2 and local T3 production, which is essential for iCAF maturation. In turn, iCAF further expressed D2 amplifying T3 signaling and sustaining a feed-forward loop between the epithelial and mesenchymal compartments. In this scenario, TH action shapes the tumor-supportive niche, facilitating tumor growth and maintenance.

However, this study has limitations. Notably, D2 expression was assessed in only a limited number of human thyroid cancers, with no detectable expression in the sole ATC sample analyzed (data not shown). While D2 expression was high in the two PDTCs tested in our experiments—supporting findings from mouse and cell line models— in silico analysis of publicly available larger series revealed increased D2 expression in both PDTCs and a subset of ATCs (37) and Supplemental Fig. 4, confirming that our findings might also apply to the subset of ATCs with a high D2 expression. Nonetheless, the limited sample size precludes definitive conclusions about the clinical features of D2-expressing tumors. Furthermore, the molecular mechanisms through which TH promotes tumor progression remain poorly understood and require further investigation.

In conclusion, this work provides strong rationale for targeting D2-dependent TH signaling in PDTC and ATC. Our data suggest that interfering with local T3 production—via D2 inhibition or modulation of TH concentration—may impair thyroid tumor growth and attenuate the tumor-supportive function of CAFs. These findings are corroborated by data from human PDTC-derived organoid models, in which D2-positive tumors exhibited TH-dependent growth. Importantly, these results raise questions about the current practice of TSH-suppressive therapy in PDTC/ATC patients. While TSH suppression remains standard approach in well differentiated thyroid cancers to limit TSH-mediated tumor growth, its relevance in poorly and in undifferentiated tumors with active TH signaling is uncertain. A better understanding of D2 expression patterns and TH sensitivity in advanced thyroid cancers may offer new avenues for personalized therapeutic strategies.

## Acknowledgments

This work was funded by AIRC Individual Grant 2022 (Project no. 27729) to DS. MADS was a recipient for IBSA Foundation Fellowship 2023.

## Authors’ contributions

MADS wrote the original draft and performed most of the experiments. CL and TP contributed to the *in vitro* studies. CC carried out IHC and mRNA analyses; CM and SS provided histological samples for the organoid studies, VC assisted with the FACS analysis, AMC and GT conducted histological evaluations and analyses; DS and MADS contributed to the investigation and data curation. MS and DS contributed conceptual insights and critically edited the manuscript; DS conceptualized the study and supervised the project. All authors edited and reviewed the final manuscript.

## Materials and Methods

### Animals

Animals were housed and maintained in the animal facility at CEINGE Biotecnologie Avanzate, Naples, Italy. Tg: B6N.Cg (Pdgfrα-cre/ERT)467Dbe/J, Pdgfrα^tm11(EGFP)Sor/^J and nude mice (NU/NU-CD1) were purchased from Charles River Stock No 18280 and 086 respectively. LSL-BrafV600E/TPO-Cre/eYFP/TP53^lox-lox^ (ATC) mice were a generous gift from Dr. Fagin at the Memorial Sloan Kettering Cancer Center in New York. These mice express endogenous levels of the mutated BRAF oncoprotein in thyroid follicular cells at E14.5, when Cre recombinase is expressed downstream of the thyroid peroxidase (TPO) gene promoter. Dio2^lox-lox^were generated in our lab (38) and crossed with the Pdgfrα-cre/ERT as described (27). Both sexes were used for experiments as indicated. Animals were genotyped by PCR using tail DNA.

### Animal study approval

Experiments and animal care were conducted in accordance with institutional guidelines. All animal studies were conducted in accordance with the guidelines of the Ministero della Salute and were approved by the Institutional Animal Care and Use Committee (IACUC: 167/2015-PR and 354/2019-PR).

### Animal procedures

Tamoxifen (TAM) (Sigma Aldrich, T5648) was dissolved in corn oil (Sigma Aldrich, C8267)/10% ethanol (Carlo Erba, #4146052) at a concentration 10 mg/ml. Pdgfrα-cre/ERT/ Dio2^lox-lox^ mice were injected intraperitoneally with TAM for five consecutive days (80 mg/Kg of body weight) for experiments involving inducible CreERT2 and preCAFs isolation.

### Cell line and mouse primary cells

Human ATC cell lines (8505c) were previously described (39). 8505 cells present mutations in BRAFV600E /P53. Isolation of mesenchymal progenitors (preCAFs) was conducted using magnetic activated cell sorting (MACS). Brifely, adult hindlimb muscles were dissected, minced and incubated with a mix of Dispase II (Roche,04942078001) 3U/ml, Collagenase A (Roche, 11088793001) 100ug/ml in PBS 1X at 37°C in a water bath for 1h. The muscle suspension was successively filtered through 70μm cell strainers (Milteny, 130-098-462) and then spun at 500g for 10min at 4°C. *Re*-suspended cell samples were incubated for 20 minutes at 4° with CD45-biotin (Miltenyi Biotech, 130-124-209), CD31-biotin (Miltenyi Biotech, 130-119-662), and Anti-integrin alpha 7-biotin (Miltenyi Biotech, 130-128-938), followed by incubation with anti-biotin microbeads (Miltenyi Biotec #120–000–900). Cells were loaded on LD columns (Miltenyi Biotec #130–042–901). The flow through fraction was collected and incubated with an anti-Sca1-microbeads antibody (Miltenyi Biotec #130–106– 641). Cells were loaded on MS columns (Miltenyi Biotec # 130-042-201) to purify the preCAF cell subfraction. preCAFs were cultured in Dulbecco’s modified Eagle’s medium (DMEM, Microgem, AL007-500ML) and supplemented with 10% fetal bovine serum, FBS, (Microgem, RM10432-500ML), 1% penicillin/streptomycin (Gibco, #15070063), and 1% l-glutamine (Gibco, #25030024). Cells were incubated at 37°C in a 5% CO2 humidified incubator. TH exogenous treatment was a mix of T4 and T3 (30nM each).

### Cell spheroids

Spheroids were generated using the liquid overlay technique in 48-well plates previously coated with 1% agarose (m/v, in water). Homo-spheroids consisted of 8505 cells (2 X 10^3^) or pre-CAF cells (6 X 10^3^) and were cultured at 37 °C, 5% CO2. Hetero-spheroids were generated by co-seeding both cell types simultaneously, using the same cell numbers as above. To confirm that the two cell types were able to interact and form hetero-spheroids, we used EGFP labelled preCAF cells isolated from Pdgfrα^tm11(EGFP)Sor/^J mice. For all the other experiments, preCAF cells were isolated from Dio2^lox-lox^ (preCAF-D2WT) as control and from Pdgfrα-Cre^ERT2^; Dio2^lox-lox^ [preCAF-D2KO] as to assess the effects of D2 depletion The size of both homo-spheroids and hetero-spheroids was monitored by bright-field microscopy from days 3 to 7, and again on days 10 and 11 of culture. For molecular expression analysis, spheroids were collected 7 days after seeding.

### In vivo xenograft tumor assays

Female and male nude mice aged 6–8 weeks were used for the xenograft assays. A suspension of 8505 cells (3 x 10⁶ cells) in 100 µl Matrigel was injected subcutaneously into the hind flank of the mice. The mice were divided into two groups: a control group (untreated mice) and an rT3-treated group. Treatment with rT3, dissolved in water, began three days prior to inoculation and continued until sacrifice. After eight weeks, the mice were sacrificed using CO₂ and their tumors were removed, weighed and photographed for comparison.

Reverse-T3 (Sigma Aldrich #T0281) was administered to mice (C56BL6 mice) via drinking water at a final concentration of 2µg/mL until the sacrifice.

### Fluorescence-Activated Cell-Sorting analysis

Thyroid tumor cells were isolated by FACS from mice tumors as previously described (40). In brief, thyroid tumors were dissected and minced in ice-cold DMEM. Samples were then incubated with a mix of 1.5mg/mL collagenase A (Sigma Aldrich, C9891), 2mg/mL Dispase II (Roche, #04942078001) at 37°C for ∼30 minutes with intermittent vortexing every 10 minutes. After digestion DMEM was added to the tissue suspensions, filtered through a 70-µm cell strainer (Corning, #431751), and pelleted by centrifugation at 600 x g for 5 minutes, at 4°C to remove large tissue fragments. Before fluorescence-activated cell sorting (FACS), the final pellet was resuspended in cold DMEM and 1% penicillin-streptomycin supplemented with 2% FBS, and the cell suspension was filtered through a 30-μm strainer (BD Bioscience, #340598). FACS was performed using FACS Aria IIIu (Becton Dickinson) by gating for YFP+ fluorescence, YFP- and SCA1+ and YFP-/SCA1-cells. We pooled n= 3 mouse thyroid tumors in 3 individual experiments.

### Cross Sectional Area (CSA) of human organoids

In brief, Bright-field images of each organoid were captured in ×20 objective lenses using an inverted microscope (CellF*Olympus). The largest cross-sectional area (CSA) was calculated using CellF*Olympus Imaging Software. Areas were normalized to their respective day 0 area of treatment. Approximately >20 organoid/ spheroids were analyses per different conditions.

### RT- Quantitative PCR

Total RNA was extracted from Thyroid tissues using TRIzol reagent (Life Technologies) according to the manufacturer’s instructions, whereas total RNA from PDO or isolated cells using a Qiagen RNeasy Micro Kit according to the manufacturer’s instructions (Qiagen, #74004). RNAs were reverse-transcribed into cDNA by using LunaScript reverse transcriptase (New England BioLabs Inc., Ipswich, MA, USA) according to the manufacturer’s instructions. Quantitative real-time PCR was performed using iQ5 Multicolor Real Time Detector System (BioRad Laboratories) with the fluorescent double-stranded DNA-binding dye SYBR Green (Applied Biosystems). Cyclophilin A gene served as the housekeeping gene controls for ΔCT calculations [ΔCT = (CT of the target gene) − (CT of housekeeping genes)]. Fold expression values were calculated using the 2−ΔΔCT method, where ΔΔCT = (ΔCT of the treatment sample) − (ΔCT of control samples) (with the control value normalized to 1). Three technical replicates were performed for all qPCR experiments (Primers used in Table S2).

### Western blot analysis

Total protein extracts from tumor tissues were run on a 15% sulfate-polyacrylamide electrophoresis gel and transferred to an Immobilon-P transfer membrane (Millipore). The membrane was then blocked with 5% BSA in phosphate-buffered saline, probed overnight at 4°C with appropriate antibodies (see Table S3) washed and incubated with horseradish peroxidase-conjugated anti-mouse or anti-rabbit immunoglobulin G secondary antibody (1:3000). The membrane was incubated with anti-GAPDH antibodies (1:1000, Santa Cruz) as a loading control. Western blots were run in triplicate. Antibody-labeled protein bands were revealed by using the Immobilon Western Chemiluminescent HRP Substrate (Millipore, WBKLS0500). The membrane images were analyzed using Image Lab version 5.2.1 (Biorad Laboratories) software.

### Patient information and sample collection

The patients were enrolled at the Department of Endocrine Surgery, Local Health Authority Naples 1 Center, Naples, Italy. This study was conducted in accordance with the Declaration of Helsinki and was approved by the Ethical Committee of University Federico II Hospital “Comitato Etico Campania ASL3” (protocol number 218/2024). Patients and tumors characteristics are summarized in Table S1.

### Establishment of PDO and primary tumor cells culture from surgical human samples

In brief, thyroid tumors were surgically removed from 4 consecutive advanced thyroid cancer patients at our Institution. The tumor samples were first mechanically and then enzymatically digested with a mixture of 0.1% collagenase A (Sigma Aldrich, C9891), 0.2% Dispase II (Roche, #04942078001) for 30 minutes at 37°C in a water bath. After digestion, DMEM was added to the thyroid tumor suspension and filtered, through a 70-μm cell strainer (Corning, #431751), and then centrifuged at 50g for 10 minutes at 4°C to remove large tissue fragments. The supernatant was discarded, and the pellet was re-suspended in DMEM and divided in two parts to obtain both primary cell culture and organoid. The primary cells were cultured in a 1:1 mixture of DMEM/F12 and supplemented with 10% fetal bovine serum 1% penicillin/streptomycin and 1% l-glutamine. Cells were incubated at 37°C in a 5% CO2 humidified incubator. The organoid were plated in 96-well plate containing 3D RGF-BME (Basement Membrane Extract Reduced Growth Factor, Corning #356231) and cultured in a medium consisting of a 1:1 mixture of DMEM/F12 supplemented with 2% BME, 10nM hydrocortisone (Sigma Aldrich, #H-0135), 1μg/mL Insulin, 5μg/mL Transferrin and 10ng/mL and 10ng/mL somatostatin (Sigma Aldrich, #S-9129) for 4 days. The organoid culture medium was replaced with the conditional medium (primary cell culture) every four days (41).

### ImmunoWluorescence and Histology

For immunofluorescent staining, cells were fixed with 4% formaldehyde and permeabilized in 0.1% Triton X-100, then blocked with 20% goat serum and incubated with primary antibody. Human thyroids tumors were collected and placed in formalin immediately and then embedded in paraffin. The samples were cut out by microtome at 10μm thickness, and stained with hematoxylin and eosin stain (H&E) (Sigma, GHS116 Hematoxylin, HT110216 Eosin), or immunofluorescence using standard protocols. Images were acquired with the ZEISS Cell Observer® fluorescence microscope and a ZEISS AxioCam MRm camera at 20x and 40x resolution.

### ImmunoWluorescence 3D

The organoids were removed from the BME and fixed in 4% paraformaldehyde (PFA) at 4 °C for 45 minutes. Then, each well containing the organoids was filled with 10 ml of cold (4 °C) PBS containing 0.1% Tween and incubated at 4 °C for 10 minutes. The organoids were then blocked in a solution of 0.2% BSA, 0.1% Triton X-100 and PBX-1X (OWB, Organoid Washing Buffer) at 4 °C for 15 minutes. Primary antibodies were then added, after which the organoids were incubated overnight at 4 °C with gentle shaking (60 rpm on a horizontal shaker). The OWB was then removed, replaced with fresh OWB and the organoids incubated for two hours with gentle shaking. After removing as much OWB as possible, a fructose-glycerol clearing solution was added at room temperature for 20 minutes. A 1 × 2 cm rectangle was drawn in the center of a slide using a PAP pen and the organoids were placed inside it. Finally, the slide was mounted with a coverslip (42). Single organoids were acquired at 20X magnification with image stacks spanning the complete organoid depth with an LSM 980 confocal system equipped with ZEN software (Carl Zeiss). Organoids area and volume were calculated by ZEISS arivis Pro software (Carl Zeiss).

### RNAscope

To detect and quantify D2 human and mouse mRNA, an ISH was performed using the RNAScope Multiplex Fluorescent V2 Assay (ACD 323100) with Opal fluorophore reagents (Akoya Biosciences). Human thyroids tumors were collected and placed in formalin immediately and then embedded in paraffin. The samples were cut out by microtome at 10μm thickness. Target probes (Mm-Dio2 #479331-C, hsDIO2 #562211-C1) were applied to the sample and baked at 40°C for 2h. Opal dyes 570 (FP1488001KT) were applied at a 1:1000 to 1:750 dilution and counterstained with DAPI. Images were taken using the ZEISS Cell Observer® fluorescence microscope and a ZEISS AxioCam MRm camera at 20x and 40x resolution.

### Statistical analysis and data graphing

Significant differences were calculated using ANOVA, and t tests with P < 0.05 were considered as statistically significant. All statistics and graphics were performed using GraphPad Prism9. In all Figures, error bars represent the S.E.M. a value of P < 0.05 was considered significant (*P < 0.05; **P < 0.01; ***P < 0.001).

