## Supplemental data for "Type 2 Deiodinase in Cancer-Associated Fibroblasts is required to sustain growth of poorly and undifferentiated thyroid cancer"

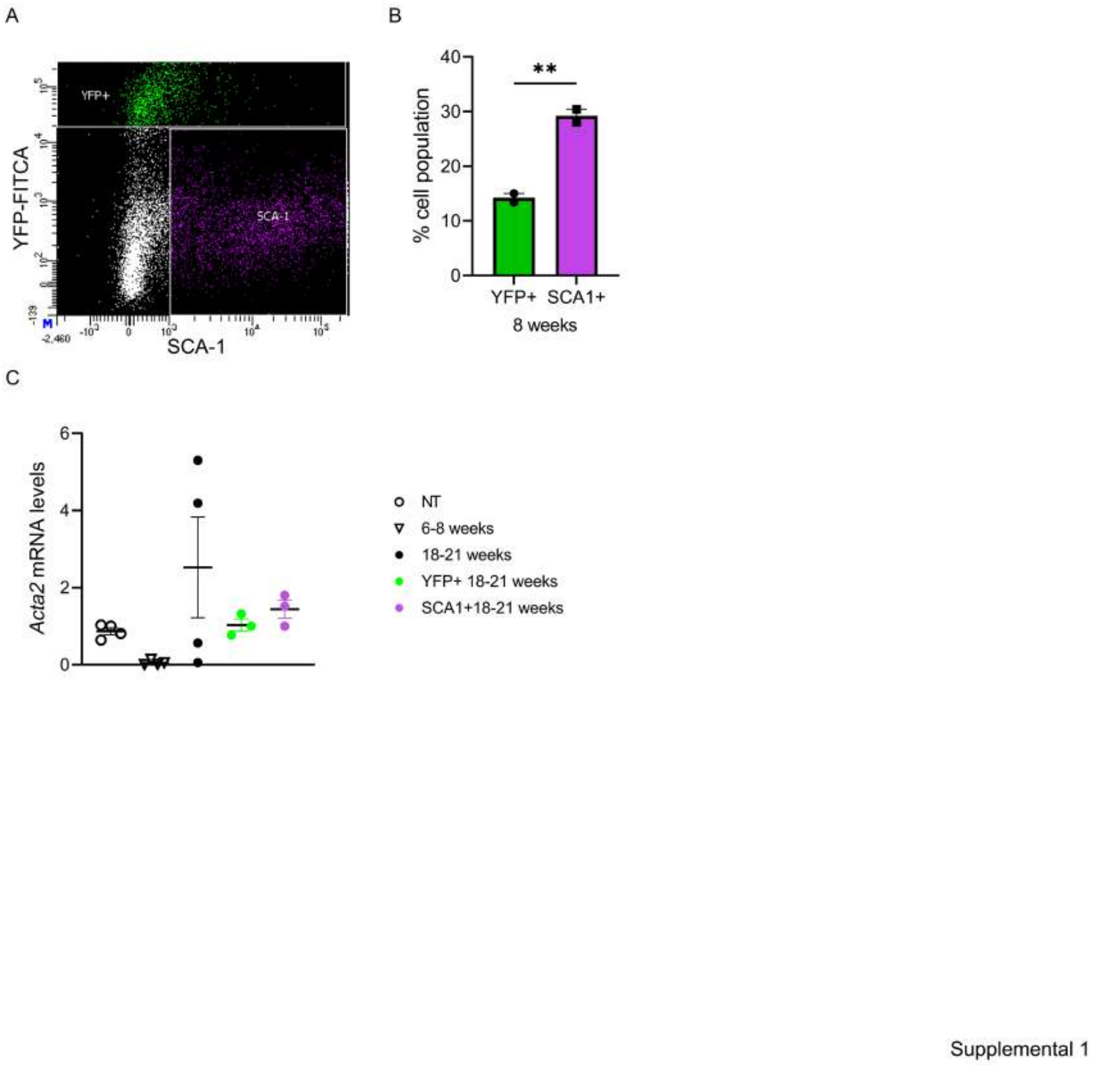

**Supplemental Fig. 1**:

(A) Representative flow cytometry dot plot showing YFP⁺ (green) and SCA1⁺ (violet) cells.

(B) Percentage of epithelial (YFP⁺) and CAF (SCA⁺) cells in pooled thyroid tumors from 8-week-old ATC mice, analyzed by FACS. Each point represents an individual experiment (pooled tumors from n=3 mice).

(C) *Acat2* mRNA expression from thyroid tumors of ATC mice at 8 and 20 weeks and YFP+ and SCA+ cells sorted from thyroid tumors of ATC mouse at 20 weeks by FACS. Each point represents individual experiment. Data are expressed as mean ± S.E.M. * $P < 0.05$, ** $P < 0.01$, ***$P < 0.0001$ using ANOVA test.

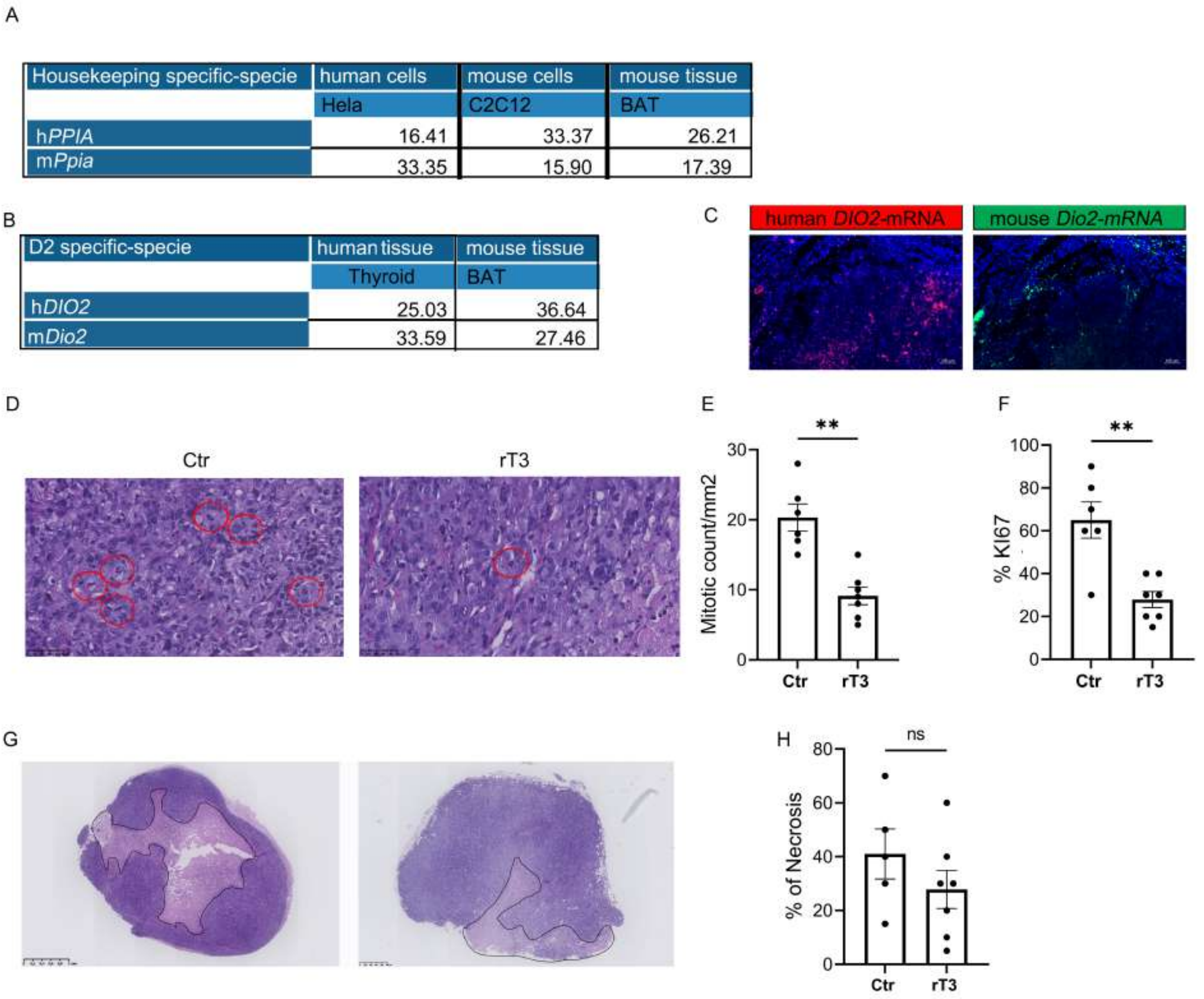

Supplemental figure 2

**Supplemental Fig. 2**:

(A) Relative expression (ΔCT values) of human h*PPIA* and mouse m*Pipia* (Housekeeping gene) in Hela, C1C12 (human and mouse cells, respectively) and BAT (mouse tissue). (B) Relative expression (ΔCT values) of human h*DIO2* and mouse m*Dio2* in Thyroid and BAT (human and mouse Tissues, respectively). (C) Fluorescent RNAscope to detect Human *DIO2* (red dots) single mRNA molecules and mouse *Dio2* (green dots) in the same cut zone of consecutive tumor sections. DAPI (blue) is used for contrast (Scale bars, 100 $\mu$m). (D) Quantification of Ki67 positive cells relative to total number of cells (Ctr=6 mice; rT3-treated=7mice). (E) H&E analysis of tumor from control and rT3-treated mice, with mitoses shown by red circles (Scale bar, 50 $\mu$m). (F) Quantification of mitotic count per mm$^2$ in tumor from control and rT3-treated mice. (G) H&E analysis of tumor from control and rT3-treated mice, with areas of necrosis marked in black. (H) Quantification of percentage of necrosis in tumors from control and rT3-treated mice. Each point represents an individual. Data are expressed as mean ± S.E.M. * $P < 0.05$, ** $P < 0.01$, ***$P < 0.0001$ using t test.

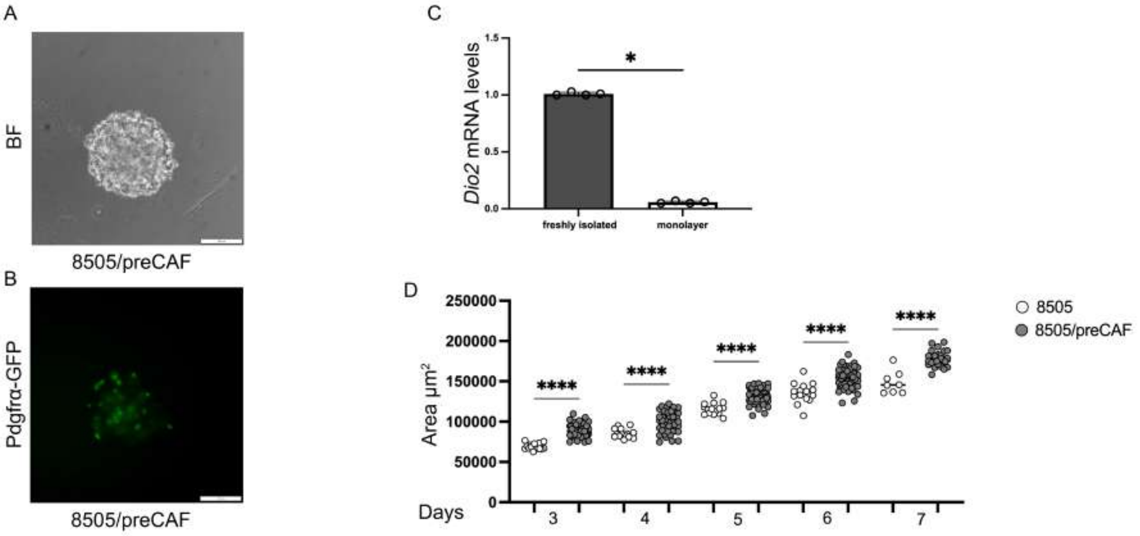

Supplemental Figure 3

**Supplemental Fig. 3:**

(A-B) Representative brightfield (BF) images (A) and corresponding epifluorescence for Pdgfrα-GFP that marked the preCAF (B) images of hetero-type spheroids (8505/preCAF;) 7 days post-seeding (Scale bars, 100 $\mu$m).

C) Dio2 mRNA expression in freshly isolated preCAF compared to preCAF cultured as a monolayer.

D) Time-course analysis of spheroid area for mono-type 8505 and hetero-type 8505/preCAF spheroids cultured for 3 and 7 days. Data are expressed as mean ± S.E.M. from at least three independent experiments. * $P < 0.05$, ** $P < 0.01$, ***$P < 0.0001$ using ANOVA test.

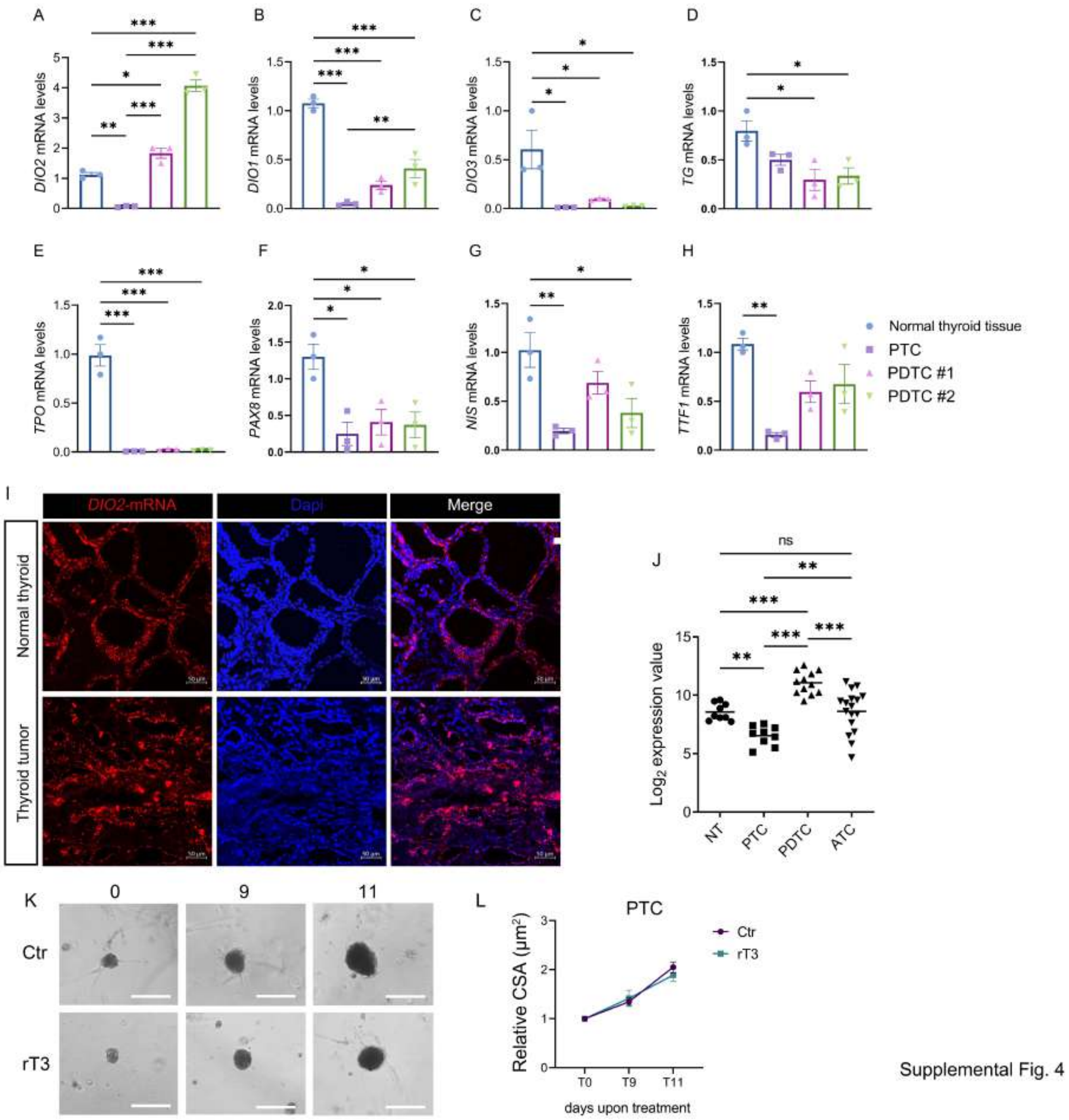

**Supplemental Fig. 4:**

**(**A-C) Deiodinases mRNA expression, *DIO2* (A), *DIO1* (B) and *DIO3* (C). Each point represents a technical replicate. D-H Thyroid-specific markers mRNA expression *TG* (D), *TPO* (E), *PAX8* (F), *NIS* (G) and *TTF1*(H) Each point represents a technical replicate.

I) Fluorescent RNAscope to detect *DIO2* single mRNA molecules (red dots). DAPI (blue) is used for contrast on section from thyroid from patient PDTC #2 (Scale bars, 50 $\mu$m). J) Array analysis of type 2 deiodinase gene (*DIO2*) expression in human thyroid cancer from the series GSE3467 and GSE76039 (Landa *et al.* 2016) analyzed using Expression Console. (K-L) Representative BF images of patient-derived organoids (patients PTC (K) and relative cross-sectional area (L), upon treatment with rT3 at different time points (Scale bar, 200 μm), areas were normalized to their respective day 0 area of treatment.

Data are expressed as mean ± S.E.M. of at least three independent technical replicates. * $P < 0.05$, ** $P < 0.01$, ***$P < 0.0001$ using ANOVA test.

### Tables

**Table S1**. Patient information and histological and mutational data of thyroid tumors.

| *Patient* | *Age at diagnosis* | *Sex* | *Histology* | *TNM* | *Sample* | *Driver gene mutations* |
|---|---|---|---|---|---|---|
| *PTC* | 78 | Male | Tall cell PTC | pT4b pN1b M1 R2 | Primary tumor | *BRAF* V600E |
| *PDTC #1* | 59 | Male | PDTC | pT3a pN1b Mx R0 | Primary tumor | NA |
| *PDTC #2* | 72 | Male | PDTC | pT3a pN1b M1 R0 | Primary tumor | PIK3CA |

Abbreviation list: PTC, papillary thyroid carcinoma, PDTC, poorly differentiated thyroid carcinoma, NA, not available.

**Table S2**

| Oligonucleotides used for real-time RT-PCR | | |
|---|---|---|
| **Human Gene** | **Forward primer** | **Reverse primer** |
| *DIO2* | TGCTGATCACACTGCAAATTC | CCTCACCCAATTTCACCTGT |
| *DIO3* | CCTGGGACTCTGCTTCTGTAAC | GGGGTGTAAGAAAATGCTGTAGAG |
| *NIS* | CCCTCATCCTGAACCAAGTG | GATCCGGGAGTGGTTCTG |
| *PAX8* | AAGTCCAGCATTGCGGCACA | GAGGGAAGTGCTTATGGTCC |
| *TTF1* | GCCGTACCAGGACACCATGAG | CAGGTACTTCTGTTGCTTGAAG |
| *TG* | GTGCCAACGGCAGTGAAGT | TCTGCTGTTTCTGTAGCTGACAAA |
| *TPO* | ACCTCGACGGTGATTTGCA | CCGCCTGTCTCCGAGATG |
| *CYA (Ppia)* | AGTCCATCTATGGGGAGAAATTTG | GCCTCCACAATATTCATGCCTTC |
| **Mouse Gene** | **Forward primer** | **Reverse primer** |
| *Cyclophilin A (Ppia)* | CGCCACTGTCGCTTTTCG | AACTTTGTCTGCAAACAGCTC |
| *Dio2* | CTTCCTCCTAGATGCCTACAAAC | GGCATAATTGTTACCTGATTCAGG |
| *Dio3* | CCGCTCTCTGCTGCTTCAC | CGGATGCACAAGAAATCTAAAAGC |
| *Col1a1* | CCCTGG TCCCTCTGGAAATG | GGACCTTTGCCCCCT TCTTT |
| *Atac2 (aSMA)* | GTCATTTTCTCCCGGTTGGC | CTGACGCTGAAGTATCCGAT |
| *Interleukin 6* | CTTCCTCCTAGATGCCTACAAAC | GGCATAATTGTTACCTGATTCAGG |

**Table S3**

| Antibodies | | | | | |
|---|---|---|---|---|---|
| **Name** | **Host** | **Source** | **Cat. number** | **Dilution** | **Application** |
| GAPDH | Mouse | Elabscience | E-AB-20059 | 1:1000 | WB |
| COL1A1 | Mouse | Santa Cruz | sc-59772 | 1:500 | IF; WB |
| Alexa Fluor 488 goat anti-mouse IgG | Goat | ThermoFisher Scientific | A-11001 | 1:500 | IF |
| Alexa Fluor 594 donkey anti-mouse IgG | Donkey | ThermoFisher Scientific | A-21203 | 1:500 | IF |
| Alexa Fluor 594 donkey anti-rabbit IgG | Donkey | ThermoFisher Scientific | A-21207 | 1:500 | IF |
| aSMA | Rabbit | Abcam | ab 5694 | 1:700 | IF; WB |
| APC-rat anti mouse CD45 | Mouse | BD Biosciences | 559864 | 1:500 | FACS |
| PE-rat anti mouse integrin-alpha7 | Rat | R&D System | FAB3518A | 1:500 | FACS |
| APC-rat anti mouse CD31 | Rat | BD Biosciences | 551262 | 1:500 | FACS |
| PE cyanine7 Ly-6A/E (Sca-1) | Mouse | Fisher Scientific | 25598182 | 1:500 | FACS |
| Ki-67 | Rabbit | Abcam | Ab-16667 | 1:500 | IF; IHC |
